# Uncovering the invisible giant: Amyloid ß plaques and their proposed association with waste removal in Alzheimer-affected human hippocampus

**DOI:** 10.1101/2025.06.01.657219

**Authors:** Ruth Fabian-Fine, Abigail G. Roman, Melanie J. Winters, Kyleena J. Lathram, Carly H. Bennett, Luwago K. Kipingi, Sondre M. Brännare-Gran, Abigail E Whitley, Chloe M. Paul, Lydia M. Altman, Ian C. Carrillo, Finn M. Joyce, Lydia A. Kragh, Theodore J. McKnight, Calum J. Reding, Leaf J. D. Reiderer, Lesley J. Rivera, Hannah A. Steen, Adam L. Weaver

**Affiliations:** Department of Biology, Saint Michael’s College, Colchester, VT 05439, USA

**Author notes:** Students participating in the 2025 Developmental Biology Course-Based Undergraduate Research Experience (CURE) course.

**Keywords:** Presenilin 1, Amyloid Precursor Protein, Tanycyte, Aquaporin4, Tau protein

## Abstract

According to the prevalent ‘Amyloid Hypothesis,’ the underlying cause for neurodegeneration in Alzheimer Disease (AD) is attributed to the accumulation of misfolded Amyloid ß and tau protein in the form of extracellular sticky plaques and neurofibrillary tangles respectively. These protein accumulations are thought to be caused by impaired waste removal. In an alternative hypothesis, we have proposed the existence of an extensive glial canal system that is likely formed by myelinated aquaporin-4 (AQP4)-expressing tanycytes and removes cellular waste from the hippocampal formation. Here, we demonstrate that tanycyte-derived waste-internalizing receptacles are immunoreactive for Aß and emanate from specialized nucleus-like organelles in the following referred to as ‘tanysomes.’ Utilizing RNA-scope in situ hybridization, we demonstrate that these receptacle-forming ‘tanysomes’ express RNA for AQP4 and the Aß-related genes, amyloid precursor protein, and presenilin 1. These findings suggest that Aß is likely synthesized where receptacle formation is observed and that Aß may play an important structural role in receptacle formation. In AD-affected hippocampus excessive amounts of Aß-immunoreactive waste receptacles emerge from tanysomes and have the appearance of plaques in Aß-immunolabeled hippocampus. Moreover, we demonstrate that the same receptacle-forming organelles exhibit strong immunolabeling for hyperphosphorylated tau protein in AD-affected tissue. We postulate that both proteins may play important structural roles in waste uptake and that hypertrophic swelling of impaired tanycytes in AD-affected brain may be due to obstructions of this extensive interconnected glial canal system.

## INTRODUCTION

Over 50 million people worldwide are believed to be affected by progressive neurodegenerative diseases (Carlsson, 2008; Sathyanarayana Rao & Jagannatha Rao, 2009; Huang *et al*., 2019; Cummings *et al*., 2021; Asher & Priefer, 2022; Breitner *et al*., 2022; Tatulian, 2022; Gustavsson *et al*., 2023). One of the most prevalent forms of neurodegeneration is Alzheimer Disease (AD) that affected roughly 6.9 million USA citizens over the age of 65, thus ranking within the leading causes of death (2024). However, despite massive international research efforts, the underlying causes for AD remain elusive. Our lack of insight into the cellular processes that trigger neurodegeneration may help explain the repeated setbacks the pharmaceutical industry has encountered in clinical trials (Carlsson, 2008; Sathyanarayana Rao & Jagannatha Rao, 2009; Huang *et al*., 2019; Cummings *et al*., 2021; Asher & Priefer, 2022; Breitner *et al*., 2022; Tatulian, 2022).

Over the past several decades, the ‘amyloid hypothesis’ has been the prevalent proposition regarding the underlying causes of AD. This theory postulates that the presence of abnormal, neurotoxic ‘amyloid beta (Aβ) plaques’ and ‘neurofibrillary tau tangles’ in the brain of AD decedents may be the result of failed waste removal from the brain explaining the detrimental formation of plaques and tangles (Carlsson, 2008; Sathyanarayana Rao & Jagannatha Rao, 2009; Huang *et al*., 2019; Cummings *et al*., 2021; Asher & Priefer, 2022; Breitner *et al*., 2022; Tatulian, 2022; Gustavsson *et al*., 2023).

Based on the amyloid hypothesis, these protein aggregates obstruct intra- and extracellular spaces thereby triggering neurodegeneration. This prevailing theory led to drugs designed to clear Aβ and hyperphosphorylated tau protein from the brain using monoclonal antibodies (Carlsson, 2008; Sathyanarayana Rao & Jagannatha Rao, 2009; Huang *et al*., 2019; Cummings *et al*., 2021; Asher & Priefer, 2022; Breitner *et al*., 2022; Tatulian, 2022; Gustavsson *et al*., 2023; 2024). However, these approaches have yielded limited cognitive benefits to patients with numerous trials failing in the clinical phase (Asher & Priefer, 2022). In addition to the suffering experienced by affected individuals and their families, the monetary implications of these failed drug trials are staggering. With an estimated annual cost of US$ 300 billion in healthcare-related expenses (2024), these numbers highlight the paramount importance to re-evaluate past and current research findings and therapeutic approaches.

*Why have we been unable to illuminate the underlying causes for neurodegeneration despite (i) massive international research undertakings, (ii) advanced scientific knowledge, and (iii) increasingly sophisticated biomedical technologies?*

We have revisited cellular structure utilizing high-resolution cell imaging, RNA-scope in situ hybridization, immunohistochemistry, and correlative light- and electron microscopy to investigate where Aβ-related genes are expressed in relation to Aβ plaques. Utilizing this knowledge, we studied how plaque-formation starts and how these plaques grow in AD-affected human hippocampus.

Due to the prevalence of rodent-models in AD research, we have also investigated mouse and rat tissue. Based on the findings presented here, we propose that the answer to the above-stated question may be misidentification of cellular structure. Here, we provide robust evidence that Aβ plaques in AD affected hippocampus may not represent random accumulations of misfolded Aβ protein deposits, but likely show abnormal proliferation and swelling of waste-internalizing receptacles that are formed by a recently-discovered glial canal system (Fabian-Fine *et al*., 2024). We show that the Aβ-immunolabeled receptacles express the Aβ-related genes amyloid precursor protein (APP) and presenilin 1 (Pres1). We postulate that the functional significance of Aβ along waste receptacles is likely their structural stabilization to prevent the collapse of these membranous receptacles during waste intake.

The findings presented here provide a new perspective into underlying causes for AD neurodegeneration.

## MATERIALS & METHODS

### Brain Tissue

The human brain tissue investigated here was obtained by Dr. John DeWitt (University of Vermont Medical Center) in the context of a previously published study (Fabian-Fine *et al*., 2024). In brief, the tissue originated from autopsy examinations of four decedents at the University of Vermont Medical Center. The tissue was obtained with full consent from the next of kin for biomedical research, diagnostic, and teaching purposes, and was fixed (see below) immediately after brain removal. All processes were in accordance with Vermont State law. Tissue from two 86-year-old decedents (post-mortem intervals of 15 and 16 h) showed accumulations of amyloid β plaques and phosphorylated tau tangles, with ABC scores consistent with intermediate (A2B2C1) and high (A3B3C2) burdens of Alzheimer disease neuropathologic change. Tissue obtained from two 76- and 35-year-old decedents (45- and 14-hours post-mortem intervals) were negative for Alzheimer disease neuropathologic change.

The mouse brain originated from adult (15-month-old) female wildtype mice (Cdh5-GCaMP8 strain). The tissue was obtained freshly upon dissection of the animals and fixed in ice-cold 4% paraformaldehyde in phosphate buffered saline 0.1 M, pH 7.4. IACUC approval was granted under PROTO202200018 (University of Vermont) and IACUC-2024-001 Fabian-Fine (Saint Michael’s College). The rat brain was obtained and processed as described previously (Fabian-Fine *et al*., 2024)

### Immunohistochemistry

Immunolabeling was conducted as described previously (Fabian-Fine *et al*., 2024). The primary antibodies utilized were rabbit anti-AQP4 (BiCell #20104), Mouse anti-APP-C99; Sigma Aldrich MABN380, and mouse anti-Myelin 6-4H2 DSHB, IOWA at dilutions of 1:100 in PBS containing 10% blocking medium overnight. The secondary fluorochrome-coupled antibodies were Cy3 goat anti-rabbit, Jackson ImmunoResearch Laboratories 111-165-003, and FITC goat anti-mouse Jackson ImmunoResearch Laboratories 115-096-072. The sections were analyzed using a confocal Zeiss AxioImager MZ with Apotome.

### Fluorochrome uptake experiments in living mouse brain

To investigate whether waste swell-bodies and associated receptacles in living mouse brain internalize Cy3 fluorochromes, we have exposed freshly dissected mouse brain to these fluorochromes as described previously (Fabian-Fine *et al*., 2025).

### Immunoperoxidase stain

Immunoperoxidase stain was carried out for Aβ and Tau protein using paraffin-embedded 5 μm-thick brain sections. The detailed procedure using the Leica Bond-3 auto staining system was described previously (Fabian-Fine *et al*., 2024). The primary antibodies used were β Amyloid 1-42 (mOC 64, AbCam) and Tau AT8 (AT8, Thermo Scientific).

### Tissue preparation for electron microscopy

Brain tissue that was processed for electron microscopy was fixed in freshly prepared 4% paraformaldehyde containing 2.5% glutaraldehyde (EMS 16019) in PBS. Post-fixation, dehydration and infiltration with Araldite was according to standard protocols as described previously (Fabian-Fine *et al*., 2024).

### Ultrathin Sectioning

The araldite embedded tissue was trimmed and sectioned with an 8-mm Diatome histo-knife using a Leica Ultracut E at a thickness of 65-nm. To prevent wrinkles, we utilized wooden toothpicks that were soaked with chloroform (Electron Microscopic Sciences, #12540) and held over the sections at approximately 2 mm distance to stretch them. It is important not to touch the sections with the toothpicks as it will damage them. The sections were collected on pioloform-coated single-slot copper or nickel grids (EMS# G2010CU). Contrasting was carried out using aqueous 1.5% uranyl acetate (6 min) and Reynold’s lead citrate (5 min) according to standard protocols. Electron microscopic examination was conducted using a JOEL 1400 electron microscope operated at 80 kV.

### RNAscope in-situ RNA hybridization

We utilized *in-situ* hybridization for the detection of AQP4, Amyloid Precursor Protein and Presenilin1 gene expression as described earlier using formalin-fixed 5 µm paraffin embedded human hippocampal sections(Fabian-Fine *et al*., 2024). The complementary probes were designed by Bio-Techne.

### Controls

Overall RNA integrity was assessed using *in situ* hybridization for the low level expressed ‘housekeeping’ gene peptidylpropyl isomerase B. For negative controls we utilized the bacterial gene dapB. We routinely assessed RNA integrity (PPIB-positive and dapB-negative) for all samples.

### Luxol H&E staining of Autopsy Tissue

This staining was performed on paraffin-embedded brain sections (5-10 μm-thick) following previously described standard protocols (Fabian-Fine *et al*., 2024).

### Quantitative Analysis

To evaluate the percentage of swell-body stages I-V in both healthy and AD-affected human hippocampus, blinded tissue sections were used to avoid bias. Images of swell-bodies throughout the CA2 and 3 regions were captured and projected onto a ∼200 cm x ∼110 cm smart board. The swell-body stages were determined and recorded collectively by the participating research team. The group attempted to evaluate comparable numbers of swell-bodies (non-AD: 314; AD: 336). To achieve these comparable swell-body numbers, one additional slice was evaluated for non-AD (4 vs. 3).

Fisher’s exact test was used to evaluate whether there was a significant association between disease state and swell-body stages. Statistics were calculated and the graph was created using Prism 10.5.0 (GraphPad Software).

### Image acquisition

Images of the Luxol H&E, anti-Aβ, and anti-tau immunolabeled paraffin sections were taken using an Olympus compound microscope with digital image acquisition capabilities.

### Image Processing

Confocal images were exported from the ZEN Blue program using the ‘Image export’ function. Pictures and figures from confocal, histological, and ultrastructural preparations were created using Adobe Photoshop.

## RESULTS

### Hippocampal tanycytes form a vast network of myelinated cell processes that project into the stratum pyramidale

Ependymal tanycytes in the ventricular lining of mouse, rat and human hippocampus form a vast network of AQP4-expressing myelinated tanycyte processes that project into the hippocampal formation (Figure 1). Individual tanycyte processes form electron-lucent ‘swell-bodies,’ each containing between one and six tanycyte-associated nucleus-like organelles referred to as ‘tanysomes.’ Swell-bodies can easily be distinguished from cell-somata, due to their lack of cytoplasmic content or organelles typically found in cell bodies (Figures 1F, 2B compared to Figure 1G). However, tanysomes give rise to waste receptacles that emanate from (*i*) the tanysome, and (*ii*) surrounding myelinated tanycyte profiles (Figure 1B-G; see also below). Interestingly, the lumina of both human and mouse tanycytes contain myelinated ring structures that swell, thereby enlarging the inner lumina of tanycyte processes in areas where these ‘luminar rings’ are inflated (Figure 1I-M). Immunolabeling demonstrates the periodic spacing of these luminar rings at 3-5 µm intervals (Figure 1M). The varicose appearance of these myelinated processes is consistently observed in mouse, rat, and human brain at both the light- and electron-microscopic levels and distinguishes tanycyte profiles from neuronal axons (Figure 1S-U). As demonstrated at the ultrastructural level in rat hippocampus, tanycytes are connected through cytoplasmic canals (Figure 1O, P) The interconnected tanycyte network sends vast numbers of varicose myelinated cell processes into the hippocampal formation where they form receptacles that project into neurons and surrounding tissue (Figure 1S-U).

**Figure 1.**
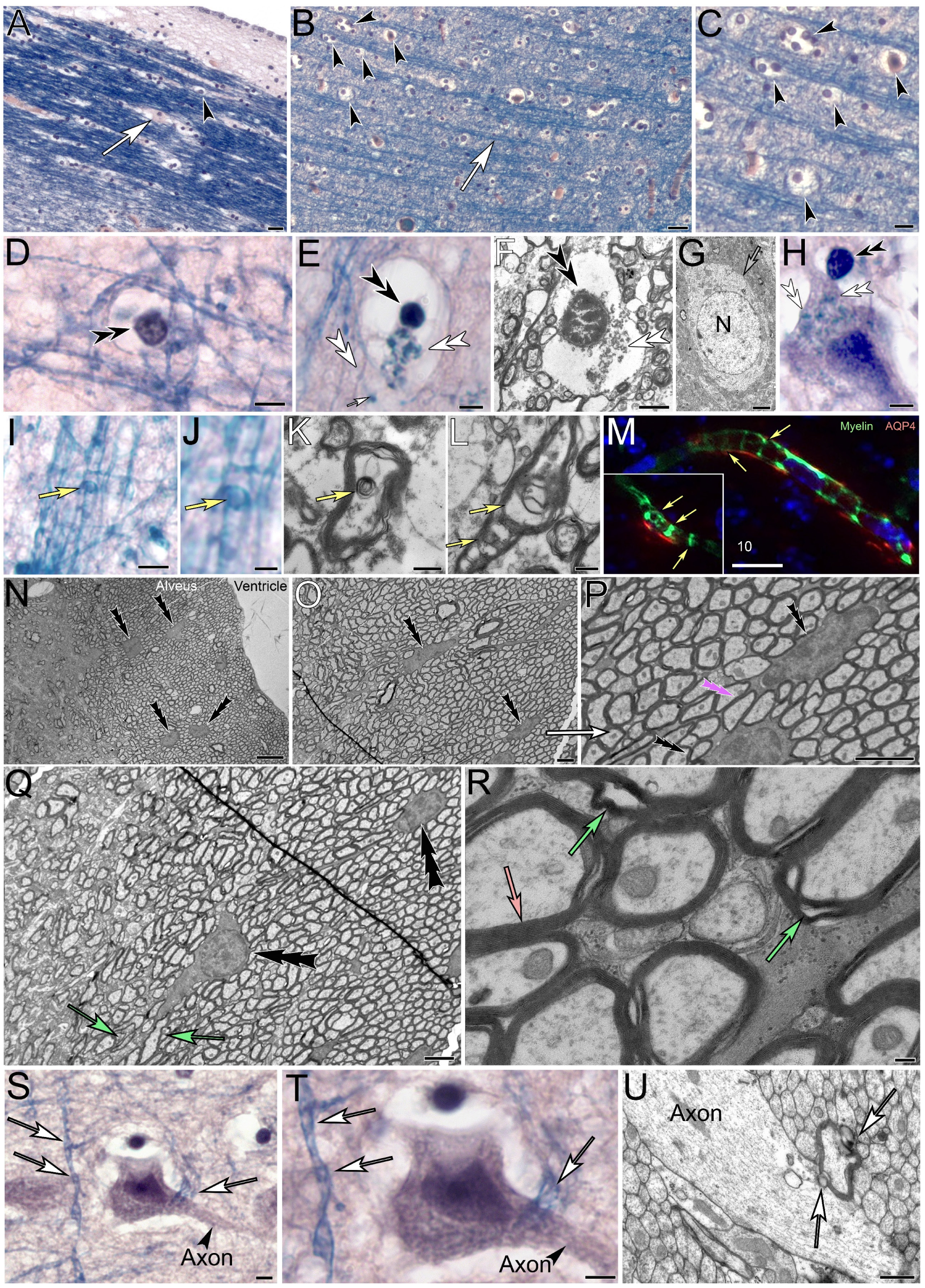
Hippocampal tanycytes form myelinated, varicose cell processes that project into the stratum pyramidale. **(A-D)** Luxol H&E-stained sections show the myelinated tanycyte processes indicated by the Luxol-blue stain (*white arrows*). Numerous swell-bodies form along the cell processes (*black arrowheads*). **(E-H)** Swell-bodies contain one or more tanysomes (*black double arrowheads*) that give rise to myelinated receptacles (*white double arrowheads; E, F, H*). The lumina of swell-bodies (*E, F*) appear translucent and lack the typical cytoplasmic organelles and structure found in cell bodies (*G, mouse hippocampal neuron*). Tanysome-derived receptacles often project into neuronal somata (*white arrowheads in H*). **(I-M)** The lumina of tanycyte processes contain myelinated ring structures (*yellow arrows H-L*) that often show increased swelling in Alzheimer-affected human brain (*K compared to L*). Immunolabeling of mouse brain for Aquaporin-4 (*M, AQP4, red*) and myelin (*green*) show similar anti-myelin immunoreactive ring structures in AQP4 immunoreactive tanycyte processes. **(N-R)** Ultrastructural depiction of the hippocampal alveus in rat brain shows somata of ependymal tanycytes (*triple arrowheads*) that give rise to vast numbers of varicosity-forming myelinated cell processes that project into the hippocampal formation. The syncytial nature of this tanycyte network is demonstrated by the cytoplasmic connections (*pink triple arrowhead in P*) formed between adjacent somata (*black triple arrowheads in P*). Please note the increasingly varicose nature of myelinated processes that exit the alveus and project into the hippocampal formation (green arrows in Q). Higher magnification shows the myelinated nature of tanycyte processes within the alveus (*light pink arrow in R*) that contain varicosities (*green arrows in R*). **(S-U)** Neuron in human AD-affected hippocampus shows varicose tanycyte processes (*S, T; white arrows*) that are clearly distinct from the neuronal axon. Please note the tanycyte process that projects into the neuron near the initial axon segment. (U) Similar observations of myelinated, varicose cell profiles transecting into neuronal axons can be observed in rat hippocampus (*white arrows*)**. *Scale bars:*** *(A, B)* 20 µm*; (C)* 10 µm*; (D, E)* 5 µm*; (F, G)* 2 µm*; (H, I)* 5 µm *(J)* 2.5 µm*; (K, L) 0.5* µm*; (M) 10* µm*; (N) 5* µm*; (O-Q) 2* µm; *(R)* 100 nm*; (S, T)* 5 µm *(U)* 0.5 µm.

**Figure 2.**
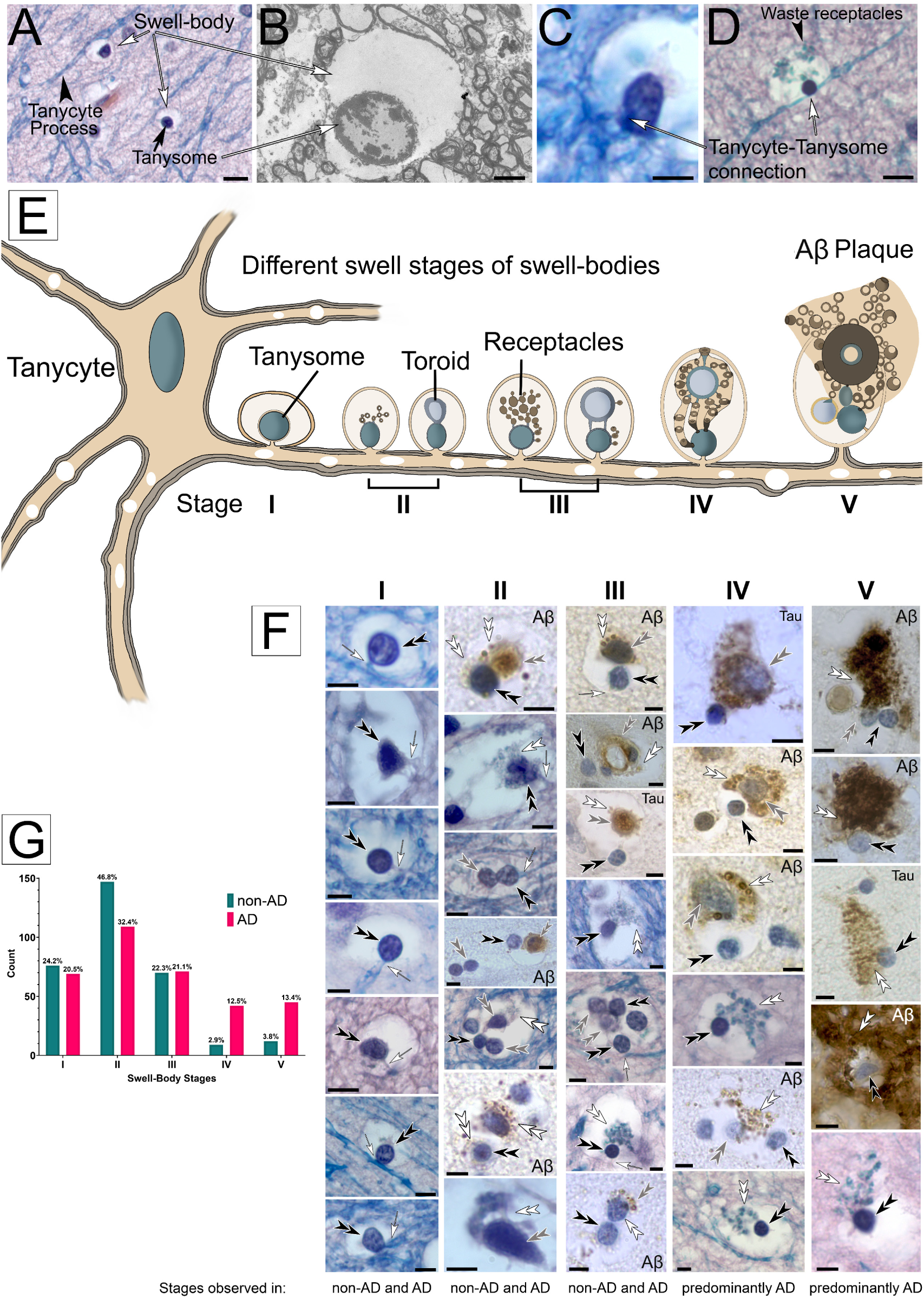
Histological, anatomical and immunohistochemical characterization of tanycyte-derived swell-body stages in human hippocampus. **(A)** Electron-lucent swell-bodies containing nucleus-like tanysomes form along varicose Luxol-blue stained tanycyte processes in Luxol H&E-stained preparations. **(B)** The electron-lucent nature of the tanysome lumen is apparent at the ultrastructural level. **(C, D)** Luxol H&E-stained swell-bodies with associated tanysomes. Please note the physical connection between tanysome and tanycyte processes (white arrows). Black arrowhead: Luxol-stained receptacles within swell-body. **(E)** Schematic drawing of tanycyte with swell-bodies that show different stages of receptacle formation (Stage I-V). For clarity, only one tanycyte process with swell-bodies is drawn. **(F)** Histological and immunohistochemical images corresponding to the schematic stages shown above each vertical image column. Stage I swell-bodies show predominantly clear lumina that contain tanycyte-associated tanysomes; Stage II swell-bodies contain tanysomes that give rise to receptacles or larger toroids (schematic drawing left and right, respectively). Please note the anti-Aβ and anti-tau immunolabeled nature of the toroids and receptacles. In Stage III swell-bodies, both receptacles and toroids swell and detach from the tanysomes, while remaining connected through slender cell processes. Swelling toroids form receptacles around their periphery that are strongly Aβ-immunolabeled. This swelling increases in Stage IV. In Stage V, both toroids and receptacles project out of the swell-bodies into the surrounding tissue or adjacent cells. Both toroids and receptacles show increasingly strong anti-Aβ and anti-tau immunolabeling. White double arrowheads: receptacles; black double arrowheads: tanysomes; grey double arrowheads: toroids; white arrows: connective cell processes between tanysome and tanycytes. All histological images are stained for Luxol-H&E. **(G)** The results of Fisher’s exact test (p < 0.0001) indicate a significant association between disease state and swell-body stages. Percentages above the bars are calculated from the total number of swell-bodies within a disease state (non-AD: 314; AD: 336). Scale bars: (A) 10 µm; (B) 2 µm; (C) 5 µm; (D) 10 µm; (F) 5 µm. Schematic not drawn to scale.

#### Tanycyte-derived swell-bodies differentiate waste-internalizing toroids and receptacles that show strong immunolabeling for anti-Aβ and anti-tau protein

Light- and electron-microscopic investigation of swell-bodies in human, non-AD affected hippocampus shows that each swell-body contains one or more nucleus-like organelles that stain for nuclear stain and are referred to as ‘tanysomes’ (Figure 2). Three primary features distinguish tanysomes from conventional cell nuclei. *(i)* The lumina of swell-bodies are void of cytoplasm and appear electron-lucent at both light and electron-microscopic levels (Figure 2A, B). *(ii)* tanysomes are penetrated by myelinated tanycyte processes (Figure 2C, D) through clearly visible pores that can reach diameters of >2 µm (Figure 2C, see also below). *(iii)* tanysomes give rise to myelin-derived receptacles that internalize electron-dense material consistent with cellular waste (Figures 2; 3; see also below). We have identified five distinct stages of receptacle differentiation in swell-bodies investigating both non-AD and AD-affected brain tissue. *Stage I* swell-bodies are ∼6-20 µm in diameter, have electron-lucent lumina and contain one tanycyte-associated tanysome (Figure 2F, *Stage I*). *Stage II* tanysomes give rise to either donut-shaped ‘toroids,’ several smaller receptacles or both. The receptacles stain for Luxol-blue, a histological marker for myelin. Interestingly, both toroids and receptacles also show immunolabeling for anti-Aβ protein in both AD and non-AD affected hippocampus. In AD-affected tissue, these structures are also labeled for anti-tau protein (Figure 2F, *Stage II*). In *Stage III* swell-bodies, the toroids swell and separate from the tanysomes but remain connected to the tanysomes via thin translucent connections. Increasing numbers of waste receptacles can be observed in individual swell-bodies (Figure 2F, *Stage III*). Both toroids and receptacles continue to increase in both size and number in Stage *IV* swell-bodies (Figure 2F, *Stage IV*) and project out of the swell-bodies in Stage *V*. The anti-tau and anti-Aβ immunolabeling observed in AD-affected tissue appears more prominent in Stage *V* compared to Stages I-IV and is consistently associated with both receptacles and toroidal structures (Figure 2F). The results of Fisher’s exact test *(p* < 0.0001) indicate a significant association between disease state and swell-body stages (Figure 2G).

### Tanysomes and associated receptacles and toroids express the Aβ-related genes amyloid precursor protein (APP) and presenilin 1 (Pres1) and give rise to Aβ-plaques

Immunolabeling of AD-affected human hippocampus for Aβ, anti-Tau protein, the Aβ-related protein APP in addition to the AQP4 water channel demonstrate the association of these proteins with toroids and waste receptacles in swell-bodies (Figure 3). The appearance of immunolabeled swell-bodies and receptacles is consistent in size, shape, and location with Luxol blue H&E-stained preparations (Figure 2). Furthermore, gene expression patterns for APP, Pres1, and AQP4 are consistent with the expression of these proteins in swell-bodies (Figure 3l-O; Figure 4A-H) and support our hypothesis that Aβ and AQP4 may play important functional roles in waste uptake into tanysome-derived waste receptacles (see discussion). Immunolabeling for Aβ (Figure 3P), AQP4, and APP (Figure 3Q-S) reveals a vast network of smaller receptacles that emanate from swell-bodies and tanycyte processes (see also below). These smaller immunolabeled sites may easily be dismissed as background labeling. However careful examination of brain sections using confocal microscopy and Differential Interference Contrast microscopy consistently show that these smaller immunolabeled sites that surround swell-bodies depict numerous translucent receptacles whose circular centers are stained for AQP4, APP, and Aβ (Figure 3P-S).

**Figure 3.**
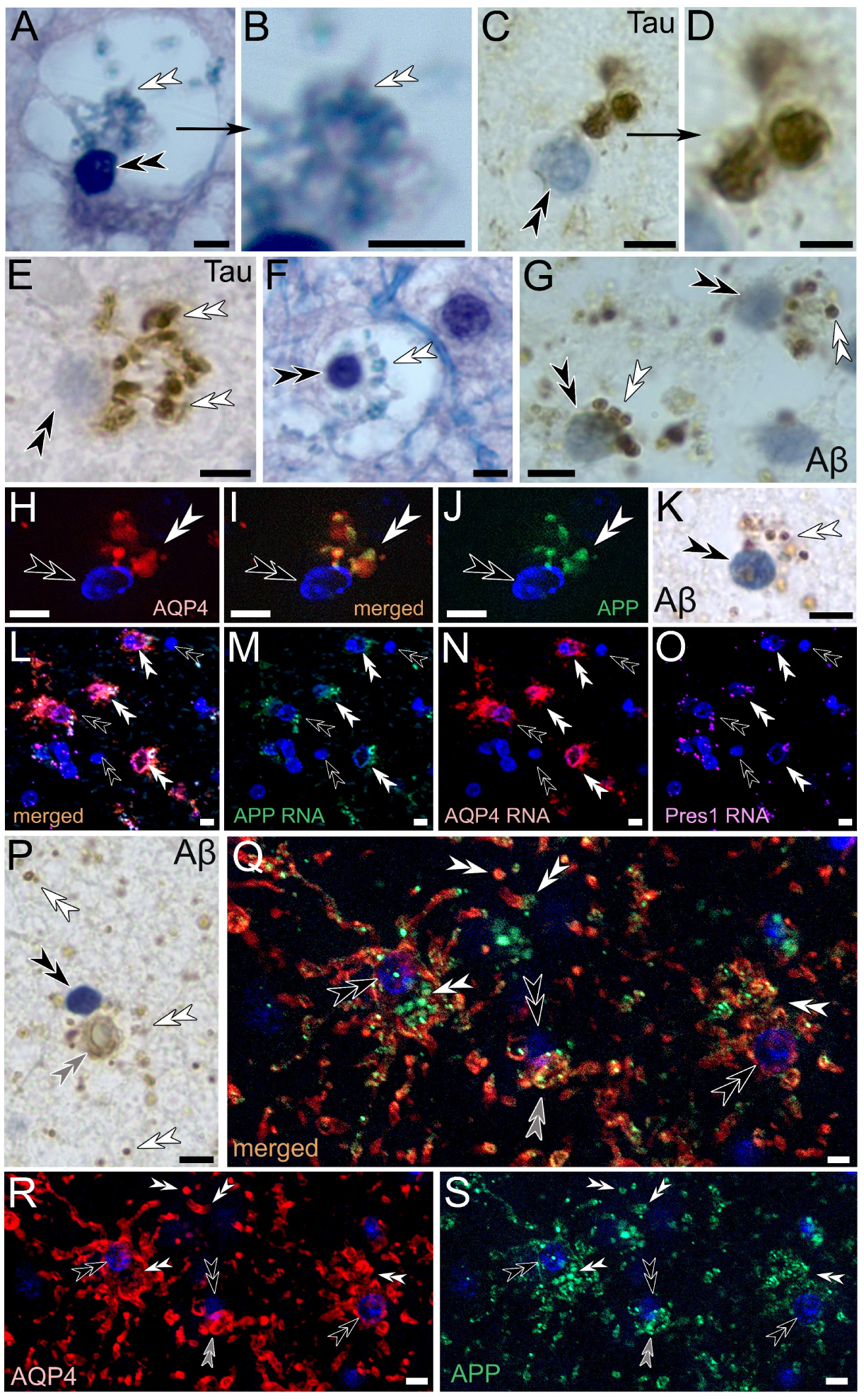
Immunohistochemical characterization of human tanysome-associated waste receptacles in swell-bodies. **(A-G)** Receptacles emanating from tanysomes stain for Luxol blue indicative of myelin (*A, higher zoom in B, F*), anti-hyperphosphorylated tau protein (*C, higher zoom in D, E*), and anti-Aβ (*G*). **(H-O)** Immunolabeling for aquaporin-4 (*AQP4, H, I*) and amyloid precursor protein (*APP, I, J*) labels receptacles that emanate from tanysomes (*Hoechst blue nuclear stain*). The observed APP-immunolabeling is consistent with the anti-Aβ immunolabeling observed on tanysome-derived receptacles (*K*). **(L-O)** RNA-gene expression for APP (*L, M*), AQP4 (*L, N*) and Presenilin1 (*Pres1, L, O*) observed along tanysomes and associated toroids and receptacles are consistent with the proposed formation of Aβ-lined AQP4 expressing receptacles in swell-bodies. **(P-S)** Tanysome-associated receptacles show immunolabeling for Aβ (*P*) and APP (*green*, *Q, S*) in AQP4-immunoreactive (*red, Q, R*) swell-bodies. Please note restriction of immunoreactivity to tanycyte processes and receptacles and the presence of numerous immunoreactive receptacles outside of swell-bodies that is consistently observed in both Aβ and AQP4/APP immunolabeled human brain of both AD and non-AD affected hippocampus (*white double arrowheads P, Q*). *White double arrowheads*: receptacles; *black double arrowheads:* tanysomes*; grey double arrowheads:* toroids. **Scale bars:** *(A)* 5 µm*; (B) 2* µm*; (C) 5* µm*; (D-E) 2* µm*; (F-S) 5* µm.

**Figure 4.**
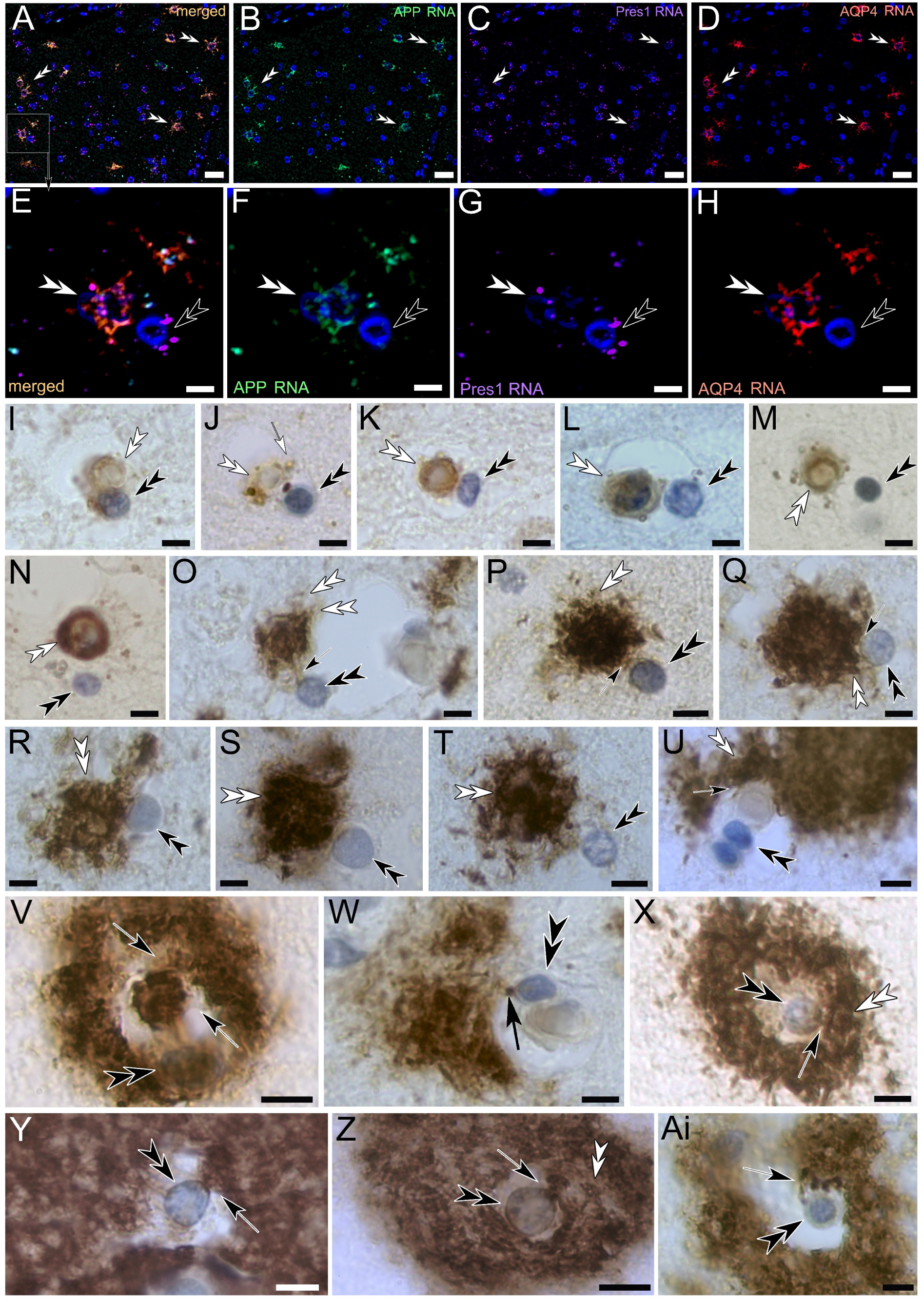
Progressive formation of Amyloid beta plaques in Alzheimer affected human hippocampus. **(A-H)** Swell-bodies in human hippocampus show RNA-expression for amyloid precursor protein (*APP, green*), presenilin 1 (*Pres1, magenta*) and aquaporin-4 (AQP4, red); *Blue signal*: DAPI nuclear stain, higher zoom shows the gene-expression along tanysomes (*black double arrowheads*) and associated toroids and receptacles (*white double arrowheads)*. **(I-M)** These findings are consistent with the modestly Aβ-immunolabeled nature of swell-body derived toroids and receptacles in AD-unaffected brain tissue that is void of Aβ-plaques. **(N-T)** In AD-affected brain tissue, both receptacles and toroids start to proliferate and swell. **(U-Ai)** Investigation of larger AB plaques shows the emergence of numerous Aβ-immunolabeled waste receptacles from tanysomes and associated toroids (*black arrows*). **Scale bars:** (A-D) 15 µm; (E-Ai) 5 µm.

### Progressive formation of Aβ-plaques

Figure 4I-Ai demonstrates the progression from Stage II swell-bodies to extremely hypertrophic Stage V swell-bodies in AD-affected Aβ-immunolabeled human brain. *Stage II* swell-bodies (Figure 4I) progressively transition into *Stage III* where the toroids and emanating receptacles separate from their associated tanysomes (Figure 4J-M). As the toroids and associated receptacles continue to swell and transition into *Stages III* and *IV*, the Aβ-immunolabeling appears more prominent (Figure 4N-Q). As these swell-bodies progress into *Stage V* (Figure 4R-Ai), the areas surrounding affected swell-bodies are densely obstructed with excessively forming Aβ-immunolabeled receptacles that emerge from tanysomes (e.g., Figure 4Y) and associated toroids (e.g., Figure 4U, V). *Stages IV* and *V* are particularly abundant in the ventricular lining. Other swell-bodies in AD-affected brain, predominantly in and near the ventricular lining, show increasingly hypertrophic swell-bodies that consist of swelling toroids with forming receptacles around their periphery (Figure 4O-T). The physical association of the Aβ-immunolabeled toroids and receptacles with tanysomes is clearly visible (Figure 4O-Q; Figure 5). Larger Aβ-plaques often form around (*i*) several adjacent swell-bodies, or (*ii*) individual swell-bodies that contain multiple tanysomes that are often located in different optical focal planes. In these plaques, large amounts of swelling receptacles can be seen adjacent to or surrounding tanysome-containing swell-bodies (Figure 4U-Ai, Figure 5 A-M). In Figure 5C, we have provided a schematic drawing of the stage V plaque depicted in Figure 5D to illustrate the formation of smaller, translucent receptacles that emerge form peripheral ‘chimney-like’ canals (corresponding arrows in 5C and D). Figure 5E shows the peripheral area of a larger plaque demonstrating the abundance of small, translucent receptacles that emerge from strongly Aβ-immunolabeled canals. The emergence of translucent waste receptacles is also demonstrated in Figure 6 where we have performed correlative ultrastructural investigations of these receptacles.

**Figure 5:**
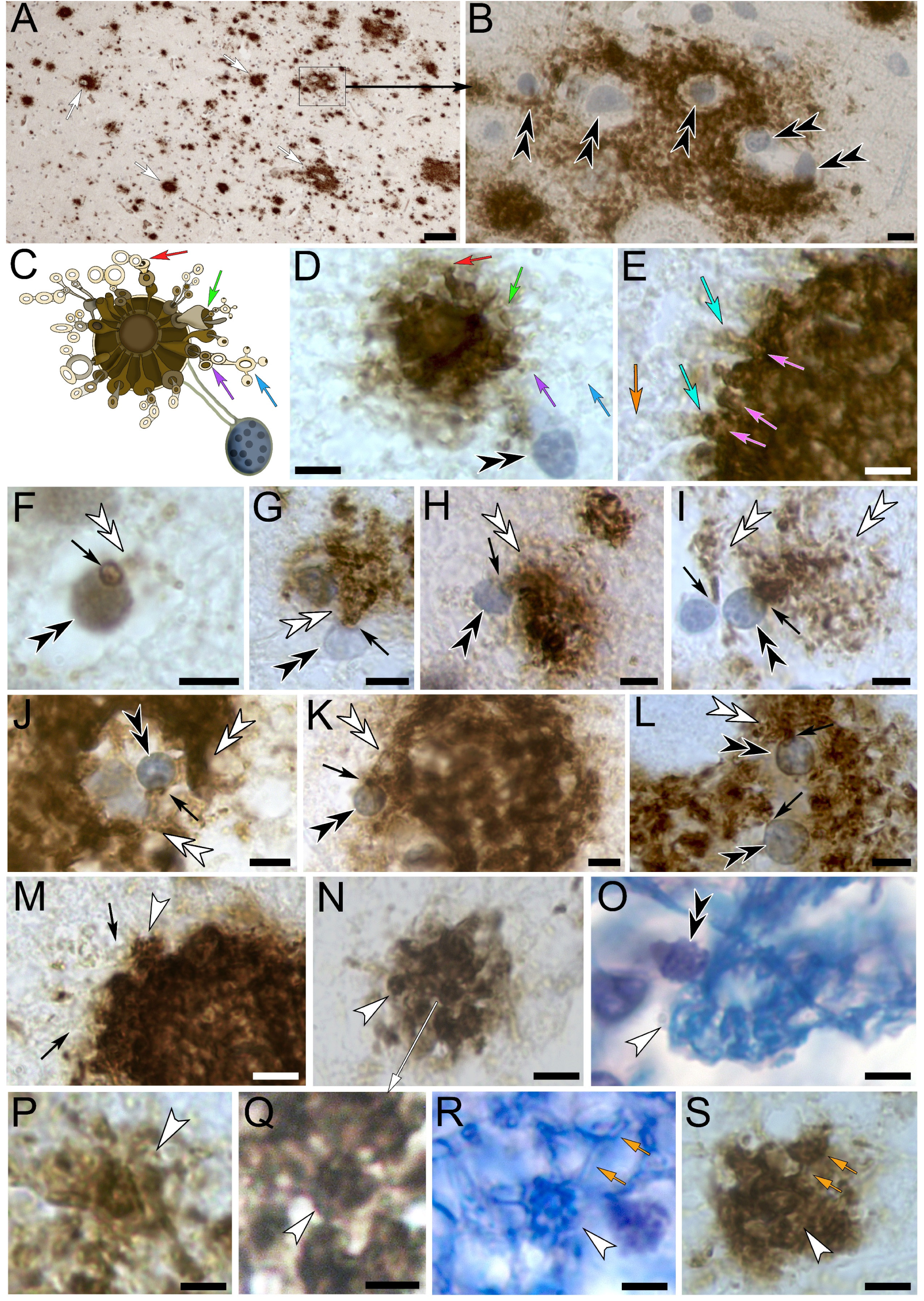
Anatomical features of Aβ plaques in AD-affected human hippocampus. **(A, B)** Dense accumulations of Aβ plaques in the hippocampal alveus (*white arrows*) are associated with tanysome-containing swell-bodies. The boxed area in *A* shown at higher magnification in *B* reveals the accumulation of tanysome-derived Aβ-immunoreactive waste receptacles (*black double arrowheads*). **(C, D)** Schematic drawing of the tanysome-associated toroid in *D* illustrates the emergence of translucent waste receptacles from peripheral canals (*colored arrows show respective areas in C and D*). Note the translucent nature of newly forming receptacles (*blue arrow in D*) **(E)** Partial image of a larger plaque shows strongly Aβ-immunolabelled ‘chimney-like’ peripheral canals (*pink arrows*) from which large numbers of forming translucent receptacles emerge (*turquoise arrows*). Please note the faint Aβ labeling of receptacles that emerge from the plaque (*turquoise arrows*) compared to the unstained nature of more distant receptacles (*orange arrow*). **(F-L)** Tanysomes (*black double arrowheads*) form Aβ-immunolabeled pores (*arrow in F*) from which large numbers of Aβ-immunolabeled toroids and receptacles emerge (*white double arrowheads*) in areas where Aβ plaques are found. **(M, N)** Toroid-shaped plaque consisting of a centrally located toroid from which peripheral receptacles emerge (*white arrowhead*). The latter in turn give rise to receptacles that appear translucent (*black arrows in M*). **(O)** Tanysome-associated toroid in Luxol H&E-stained human alveus shows a centrally located toroid that contains Luxol-blue stained circular receptacles and ring structures (white arrowhead). (P-S) Higher zoom of star-shaped receptacles (*white arrowheads*) observed in Aβ-plaques and Luxol H&E-stained alveus. Radial extensions from the centrally located toroidal structure form additional receptacles (*orange arrows*). Image in *Q* is a higher zoom of the plaque shown in image *N. **Scale bars:*** (*A*) 15 µm; (*B, D-O*) 5 µm; (*P-S*) 2.5 µm. *Schematic not drawn to scale*.

**Figure 6:**
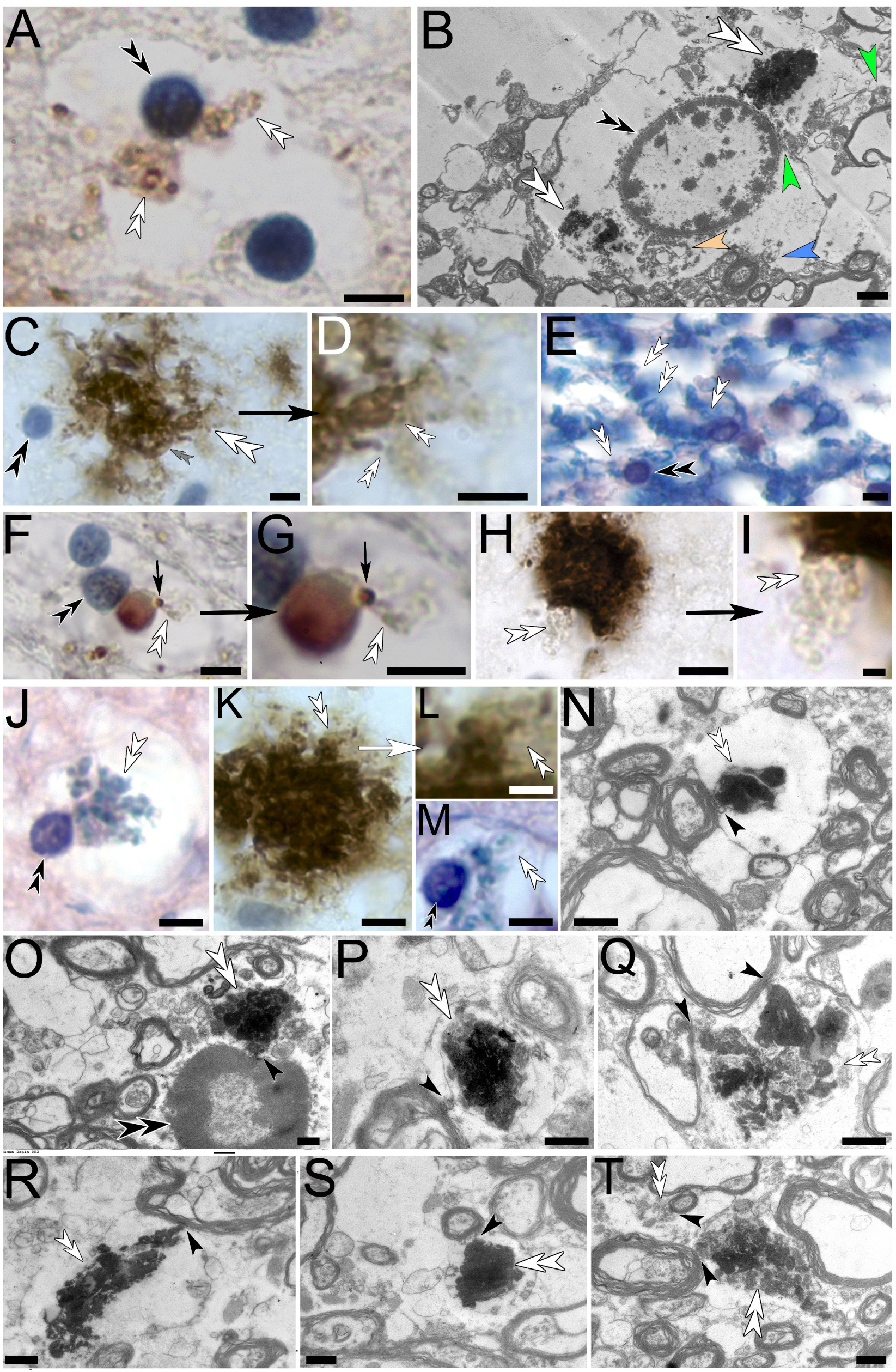
Correlative light and electron-microscopic characterization of tanysome-derived waste receptacles in AD-affected human hippocampus. **(A)** Amyloid β-immunolabeled swell-body with receptacle-forming tanysome. Please note the strong circular immunolabeling in the center of forming receptacles. **(B)** Ultrastructural depiction of a swell-body with centrally located tanysome and waste receptacles. Some receptacles are filled with electron-dense material (white arrowheads) while others appear less electron-dense (beige arrowhead). Receptacles also emanate from myelinated tanycyte profiles that border the outer periphery of the swell-body (blue arrowhead) and can be seen to form connections to both tanysomes and surrounding myelinated cell profiles (green arrowheads).**(C, D)** Tanysome with associated Aβ-immunolabeled toroid (outlined by grey double arrowheads) from which lateral strands of waste receptacles emanate (white double arrowhead, higher zoom in D). **(E)** Luxol H&E-stained tanycytes in the alveus show Luxol-stained receptacles emanating from tanysomes. **(F-I)** Amyloid β-immunolabeled toroids from which translucent waste receptacles emanate (black arrows, G and I depict higher zooms of images F and H respectively). Please note the strong immunolabeling of the pore from which receptacles emanate in F, G (white arrows). **(J, K)** Tanysome-associated toroids with emanating waste receptacles. **(L-N)** Higher zoom of peripheral waste receptacles (*L*) indicated in *K* show emanating peripheral receptacles similar to the one shown in Luxol H&E-stained swell-body. Correlative electron-microscopy depicts a myelinated receptacle associated with electron-dense material consistent with internalization of cellular waste. **(O-T)** Electron-micrographs of waste-internalizing receptacles that emanate from myelinated tanycyte profiles. White double arrowheads: receptacles; black double arrowheads: tanysomes; grey double arrowhead: toroid; black arrowhead: areas in which electron-dense waste receptacles emanate from myelinated tanycyte profiles or tanysomes. Scale bars: (A) 5 µm; (B) µm; (C-H) 5 µm; (I) 1 µm; (J, K) 5 µm; (L) 2 µm; (M) 5 µm; (N-T) 500 nm.

### Amyloid β-plaques consist of interconnected star-shaped receptacles that can also be observed in Luxol H&E

High magnification examination of individual Aβ-plaques shows that they consist of star-shaped receptacles each comprised of a central receptacle from which peripheral canal-like extension emanate that interconnect adjacent receptacles (Figure 5K-S). To demonstrate the abundance and consistency by which large amounts of Aβ-immunolabeled, and translucent receptacles emerge from tanysomes and toroids in AD-affected human hippocampus, we have included a larger than usual number of images that show this process (Figures 4-6). Ultrastructural investigations of waste receptacles in AD-affected hippocampus demonstrate the association of both tanysomes and myelinated cell profiles with strands of electron-dense receptacles consistent with the hypothesis that these receptacles internalize cellular waste (Figure 6). Our ultrastructural investigations show the association of numerous waste receptacles with myelinated profiles within and outside of swell-bodies. These observations are consistent with Luxol H&E-stained preparations that show blue coloration of forming receptacles within swell-bodies indicative of myelination (Figure 6J, M).

### Anti-Tau immunolabeling of AD-affected human hippocampus is associated with tanysomes and emanating receptacles and toroids

Investigation of anti-Tau immunolabeled AD-affected tissue demonstrates that this protein is also associated with swell-bodies and forming receptacles (Figure 7). Interestingly, the appearance of anti-Tau immunolabeled swell-bodies in human AD affected brain shows strong resemblance to swell-bodies in living mouse brain after exposure to Cy3 fluorochrome indicative of fluorochrome internalization into the swell-bodies and circular receptacles contained within the swell-bodies (Figure 7D).

**Figure 7:**
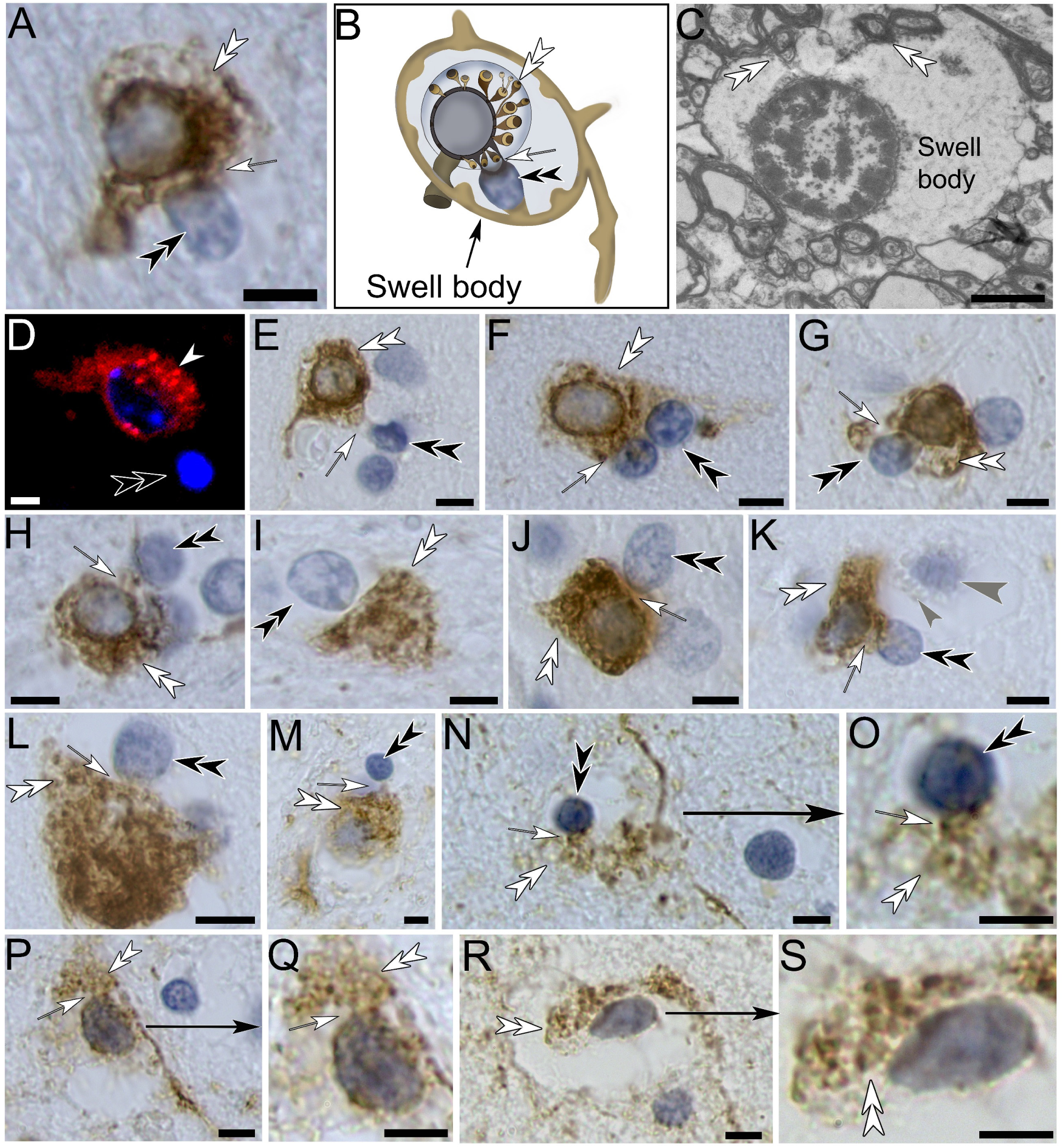
Anti-Tau immunolabeling of AD-affected human hippocampus. **(A)** Swell-body with receptacle-forming toroid shows that the anti-tau labeling is associated with receptacle formation. Schematic drawing of swell-body in *A*. **(C)** Electron-micrograph of swell-body with forming myelinated receptacles. **(D)** Swell-body in living mouse hippocampus that was exposed to CY3 fluorochrome shows internalization of the fluorochrome (white arrowhead). **(E-S)** Swell-bodies that show the consistency of tau-immunolabeling associated with receptacle formation from both toroids (E, F, G, H, I, J, M) and tanysomes (K, L N with higher zoom in O, P with higher zoom in Q, R with higher zoom in S). Please note the unlabeled swell-body with forming receptacles (small grey arrowhead) that protrude from nuclear-stained receptacle shaped structures (large grey arrowhead) in K. White double arrowheads: receptacles; black double arrowheads: tanysomes, white arrows: connection between tanysomes and receptacles or toroids. Scale bars: (A) 5 µm; 2 µm; (*D-S*) 5 µm. Schematic not drawn to scale.

## DISCUSSION

Here, we demonstrate that Aβ-plaques are formed by tanycyte-derived toroids and receptacles that emanate from tanysomes within swell-bodies. Utilizing both, immunohistochemistry, and RNA-scope *in situ* hybridization for the Aβ-related genes APP and Pres1, we demonstrate that their encoding RNA and proteins are expressed by tanysomes and in swell-bodies where receptacle formation is observed. Both APP and Pres1 play important roles in the formation of Aβ (Kepp *et al*., 2023). Thus, our findings together with the prominent anti-tau immunoreactivity observed in AD-affected swell-bodies support our postulation that both tau and Aβ proteins may play important roles in the structural and functional architecture of waste receptacles. Moreover, the observation of both AQP4-RNA expression and immunolabeling indicates the expression of this protein within swell-bodies, which provides a compelling explanation for the different swelling patterns observed in both swell-bodies and associated receptacles. Our postulation that tanysome-derived receptacles internalize cellular waste is corroborated by our correlative ultrastructural investigations that show the accumulation of electron-dense material within receptacles.

### Evidence in support of the glial canal hypothesis’

Based on the findings presented here, we postulate that Aβ-plaques show hypertrophic waste receptacles that emanate from tanycyte-associated swell-bodies. This postulation contradicts the prevalent ‘amyloid hypothesis’ that proposes that Aβ-plaques are due to the random accumulation of misfolded Aβ protein (Kepp *et al*., 2023; Behl, 2024).

We favor the recently proposed ‘glial canal hypothesis’ (Fabian-Fine *et al*., 2024) due to the following observations: (i) The tanysome-associated RNA-expression for APP, Pres1, and AQP4 within swell-bodies demonstrates that Aβ is likely formed within swell-bodies where Aβ-immunoreactivity is detected. This result contradicts the proposed random accumulation of this protein. (ii) The observed immunolabeling for the RNA-encoded proteins APP, AQP4, and Pres1 (Fabian-Fine *et al*., 2025) together with Aβ along toroids and receptacles that emanate from tanysomes is consistent with the RNA expression patterns that encode for these proteins. (iii) The swelling nature of tanycyte-derived receptacles, processes, and swell-bodies. Swelling of AQP-expressing cells is well established and would furthermore explain the electron-lucent nature and lack of cytoplasmic structure observed in swell-body lumina (Preston *et al*., 1992; Wang *et al*., 2023). This swelling would furthermore explain why neuronal somata in AD-affected brain tissue are often densely obstructed with receptacles which results in cytoplasmic depletion and cell death (Fabian-Fine *et al*., 2024). Until now, such intraneuronal receptacles have been described as ‘lipofuscin’ that has been proposed to contain toxic waste that remains in neurons indefinitely (Gray & Woulfe, 2005). Considering our findings that myelin-derived tanycytes form receptacles that project into neuronal somata [see also (Fabian-Fine *et al*., 2025)], we encourage the re-investigation of the proposed lipofuscin particles and test whether they may represent myelin-derived receptacles. This likelihood would explain the prominent intraneuronal myelin-specific Luxol-stain observed in neurons of Australian cattle dogs diagnosed with neuronal ceroid lipofuscinosis, a disease that is characterized by progressive neurodegeneration [see Figure 11 in (Schmutz *et al*., 2019)]. We postulate that the structures identified as lipofuscin depict intraneuronal tanycyte receptacles that clear waste from healthy neurons. We suggest that pathological swelling of these receptacles may be due to hypertrophic impairment of the AQP4-expressing tanycytes that obstructs neuronal somata resulting in neurodegeneration(Sauvé *et al*., 2022; Fabian-Fine *et al*., 2024; Fabian-Fine *et al*., 2025).

### Proposed functional significance of Aβ, tau protein, myelin, and AQP4 in waste uptake

#### Amyloid β and AQP4

The hydrophobic nature of both Aβ (Vugmeyster *et al*., 2016) and receptacle membranes likely favors the assembly of synthesized Aβ along forming receptacles. As previously demonstrated, Aβ-immunoreactive receptacles have been described in healthy non-AD affected neurons (Kobro-Flatmoen *et al*., 2023; Fabian-Fine *et al*., 2024; Fabian-Fine *et al*., 2025). We postulate that these receptacles represent a vast tanycyte-derived network that continuously clears cellular debris from healthy neurons and their surroundings. Consistent with our findings presented here, we postulate that this clearance is generated through an AQP4-mediated convective flow that flushes waste toward the receptacles. However, such a system requires the physical stabilization of the waste-internalizing receptacles to ensure that the openings remain open and avoid collapse. In the pulmonary system, stabilization of trachea is provided by C-shaped cartilage rings. The latter prevents tracheal collapse despite the small vacuum produced during inspiration (Lambert *et al*., 1991; Rains *et al*., 1992). Interestingly, Aβ aggregates reportedly form very stable molecules consistent with the requirement of receptacle stabilization during waste uptake (Tycko, 2015). The regularly-aligned swelling luminar rings in tanycytes shown here [see also (Fabian-Fine *et al*., 2025)] would provide stabilization of the glial-canals, and the dynamic ability to enlarge their diameters during waste uptake. The proposed AQP4-mediated swelling of these luminar rings would create an inward fluid flow into the canals that draws waste with it as proposed earlier [(Fabian-Fine *et al*., 2025), Figure 11]. In this context, the previously proposed circadian dynamic of waste removal (Hablitz *et al*., 2020) that may regulate the periodic swelling of these canals provides a compelling mechanism. It remains to be investigated whether other AQP channels may be present in this system. The concept of an AQP4-mediated convective flow in context of waste removal from the brain was previously postulated in context with astrocytes and the proposed glymphatic system (Iliff *et al*., 2012).

#### Tau Protein

An additional structural requirement for the proposed glial canal system is ‘dynamic adaptability.’ To ensure adequate waste removal, the quantitatively appropriate release of waste receptacles is imperative. Too many receptacles pose the risk of neuronal obstruction and depletion; too few receptacles may cause waste buildup within the cells. Previous investigations on spider brain have shown that neurodegeneration is preceded by pathological unraveling of adjacent myelinated glial cells. As a consequence, in areas where the glial cells unravel, we have observed excessive formation of waste-internalizing glial canals that lead to the catastrophic depletion of affected neurons (Fabian-Fine *et al*., 2023; Fabian-Fine *et al*., 2024). Interestingly, in this arachnid model system, the forming glial canals are anchored to microtubules. This observation led us to propose that the dynamic instability of microtubules is a uniquely suitable mechanism to enzymatically and dynamically control the quantitatively appropriate release of waste-internalizing glial canals. Triggering mechanisms may include mechanical or biochemical signals such as increasing hydrogen concentrations caused by lysosomal activity that produces cellular waste (Fabian-Fine *et al*., 2024). Interestingly, tau protein is well known to stabilize microtubules in the mammalian brain (Barbier *et al*., 2019). As demonstrated here, its hyperphosphorylated form is abundant in AD-affected swell-bodies in which excessive receptacle proliferation is observed. It is thus feasible that microtubules may play a similar role in controlling the quantitatively appropriate release of waste receptacles and that this mechanism may fail in AD. This idea would explain the increased formation of waste receptacles that coincides with immunolabeling for hyperphosphorylated tau protein shown here.

### Tanycytes and their proposed roles

Tanycytes are ependymal glial cells that have predominantly been described along the ventricular lining of the third and fourth ventricles and are currently subdivided into four major types based on their location alpha-1 and −2, beta-1 and −2. However, the presence of additional tanycyte types has been proposed (Pasquettaz *et al*., 2021; Fabian-Fine *et al*., 2024). These ependymal glial cells that label for glial fibrillary acid protein send long, slender processes into the hypothalamic parenchyma and are proposed to have metabolic functions (García-Cáceres *et al*., 2019). Interestingly, the formation of varicose protrusions along tanycytes that contact neurons have been observed in the hypothalamus of mice (Pasquettaz *et al*., 2021). The degradation of tanycytes in AD-affected brain has previously been reported (Sauvé *et al*., 2022). The authors propose that tanycyte impairment may play a prominent role in neurodegeneration. Based on our findings here we concur with this postulation.

### Relevance of these findings for pharmaceutical approaches

Provided our hypothesis is correct and Aβ-lined toroids and receptacles in Stage I-III swell-bodies show waste-internalizing receptacles in healthy brain tissue, these findings have significant implications for drug development. Pharmaceutical approaches that target the destruction of Aβ and tau protein with monoclonal antibodies (Sevigny *et al*., 2016; Chowdhury & Chowdhury, 2023) target this protein in healthy *Stage I-III* swell-bodies *and Stage IV*, *V* swell-bodies (Aβ-plaques). This approach thus poses the risk that remaining healthy waste removal will be impaired by antibody treatment and would provide an explanation why most clinical trials utilizing this approach have repeatedly failed (Carlsson, 2008; Huang *et al*., 2019; McDade, 2019; Tolar *et al*., 2020; Asher & Priefer, 2022; Breitner *et al*., 2022).

## CONCLUSION

This study provides novel insight into the histopathology of AD-affected brain tissue and offers a new hypothesis for the formation of Aβ plaques and neurofibrillary tau tangles. We are acutely aware that we are unable to address other important topics in this study. Among other topics this includes: (*i*) the striking resemblance of brain structure identified as astrocytes in brain disease with *Stage III-V* swell-bodies shown here (see Figure 15 in (Verkhratsky *et al*., 2023)), (*ii*) the roles of myelin and glial fibrillary acid protein in the brain, (*iii*) the possible role of tanycytes with regard to gliomas, (*iv*) possible biochemical pathways that trigger receptacle formation in swell bodies, and (*v*) underlying causes that lead to excessive receptacle formation. We have addressed the latter in previous publications where we propose a link between AD and possible viral, fungal, and bacterial infections whose impact is increasingly recognized (Muzambi *et al*., 2020). It is feasible that structural proteins of such pathogens are not sufficiently catabolized prior to uptake into the glial canal system and may lead to blockage (Fabian-Fine *et al*., 2024; Fabian-Fine *et al*., 2025). The resulting increase in intracellular turgor would explain the excessive release of receptacles from these AQP4-expressing organelles. These and other topics must be addressed in more detail in future studies. We concur with previously voiced suggestions to critically re-evaluate the ‘amyloid hypothesis’ (Kepp *et al*., 2023) and consider alternative hypotheses, one of which is presented here.

## AUTHOR CONTRIBUTIONS

Conceptualization: RFF

Funding acquisition: RFF, ALW

Methodology: RFF

Project administration: RFF

Supervision: RFF

Investigation: RFF, ALW, AGR, MJW, KJL, CHB, LKK, CMP, LMA, ICC, FMJ, LAK, TJM, CJR, LJDR, LJR, HAS

Visualization: RFF, ALW, AGR, MJW, KJL, CHB, LKK, CMP, LMA, ICC, FMJ, LAK, TJM, CJR, LJDR, LJR, HAS

Formal Analysis: RFF Resources: RFF, ALW Validation: RFF

Writing – original draft: RFF

Writing – review & editing: RFF, AGR, ALW

## Data availability

Requests for the data resources should be directed to and will be fulfilled by the lead contact, Dr. Ruth Fabian-Fine. All data including access to the original histological slides examined in context of the work reported in the text will be made available by the lead contact.

## Ethics approval statement

Processing and examination of autopsy tissue was in accordance with Vermont State law.

## Patient consent statement

Brain autopsy tissue was fully consented by next of kin

## Permission to reproduce material from other sources

Not applicable

## Clinical trial registration

Not applicable

## ACKNOWLEDGEMENTS

The authors would like to thank Dr. John DeWitt for support and access to human decedent tissue. We thank Drs. Alain Brizard, Christopher Francklyn, Mark Nelson, Mark Lubkowitz, Douglas Taatjes, and Heather Driscoll for support and helpful feedback. We thank Ann and Dr. David MacLaughlin for their donation to fund the Zeiss confocal microscope. The Electron Microscopy was conducted at the Microscopy Imaging Center at the University of Vermont (RRID: SCR_018821). We thank Natalie Cashen, Dr. Gerald Herrera, Nicole Bouffard, Kyra Lee, and Brad Vietje for technical support. Research reported in this publication was supported by an Institutional Development Award (IDeA) from the National Institute of General Medical Sciences of the National Institutes of Health under grant number P20GM103449. Its contents are solely the responsibility of the authors and do not necessarily represent the official views of NIGMS or NIH.

